# Competition and Synergy Between *Prochlorococcus* and *Synechococcus* Under Ocean Acidification Conditions

**DOI:** 10.1101/337378

**Authors:** Margaret A. Knight, J. Jeffrey Morris

## Abstract

Anthropogenic CO_2_ emissions are projected to lower the pH of the open ocean by 0.2 to 0.3 units over the next century. Laboratory experiments show that different phytoplankton taxa exhibit a wide variety of responses, with some strains having higher fitness under projected future conditions, and others being negatively impacted. Previous studies have suggested that *Prochlorococcus* and *Synechococcus*, the numerically dominant picophytoplankton in the oceans, have very different responses to elevated CO_2_ that may result in a dramatic shift in their relative abundances in future oceans. Here we show that these two genera experience faster exponential growth rates under future CO_2_ conditions, similar to most other cyanobacteria that have been studied. However, *Prochlorococcus* strains have significantly lower realized growth rates due to more extreme lag periods after exposure to fresh culture media. Surprisingly, however, *Synechococcus* was unable to outcompete *Prochlorococcus* in co-culture at elevated CO_2_. Under these conditions, *Prochlorococcus*’ poor response to elevated CO_2_ disappeared, and it showed negative frequency dependence in its relative fitness compared to *Synechococcus*, with a significant fitness advantage when it was initially rare. Moreover, both *Synechococcus* and *Prochlorococcus* had faster growth rates in co-culture with each other than either had in unialgal culture. We speculate that this negative frequency dependence is an outgrowth of reductive Black Queen evolution operating on both taxa that has resulted in a passively mutualistic relationship analogous to that connecting *Prochlorococcus* with the “helper” heterotrophic microbes in its environment.

## INTRODUCTION

Human consumption of fossil fuels has increased atmospheric CO_2_ concentration from 280 ppm in pre-industrial times to ∼ 400 ppm today, directly causing a decrease in ocean pH of approximately 0.1 unit, and models predict that, without substantial changes in human activity, ocean pH will drop a further ∼ 0.3 units by the end of the 21st century (Doney *et al.*, 2009). Ocean acidification will impact marine organisms in two potentially counteracting ways. First, increased acidity reduces the stability of calcium minerals used by many organisms to form shells and skeletons (Kleypas *et al.*, 2006) and can interfere with other key physiological processes. Second, increased concentrations of dissolved CO_2_ can stimulate physiological processes that use CO_2_ as a substrate. Since CO_2_ is the primary substrate for photosynthesis, the tension between these two countervailing forces is perhaps strongest for the ocean’s photosynthetic organisms, including the microbial phytoplankton that form the base of the oceanic food web (Mackey *et al.*, 2015). While carbon availability is unlikely to limit photosynthetic rates in the open ocean, projected year 2100 CO_2_ concentrations nevertheless stimulate growth rates in laboratory cultures of several important phytoplankton taxa (Feng *et al.*, 2008, Fu *et al.*, 2008, Hutchins *et al.*, 2007, Lefebvre *et al.*, 2012, Li *et al.*, 2012, Sobrino *et al.*, 2008, Spielmeyer & Pohnert, 2012, Sun *et al.*, 2011, Tatters *et al.*, 2013). Other experiments have shown phytoplankton growing slower at higher CO_2_ (Iglesias-Rodriguez *et al.*, 2008, Hoogstraten *et al.*, 2012, Muller *et al.*, 2012, Hennon *et al.*, 2018), and in some cases different strains of the same species have opposite response to high CO_2_, even when tested side-by-side by a single researcher (Langer *et al.*, 2009). These differential responses appear to have ecological consequences, as addition of CO_2_ (and concomitant reduction of pH) to natural seawater communities induces substantial shifts in phytoplankton community composition (Tortell *et al.*, 2002, Hare *et al.*, 2007).

The effect of pH on phytoplankton ecology is qualitatively different from the effect of temperature, the other major parameter influenced by rising CO_2_. The many varieties of phytoplankton are not physiologically interchangeable, but rather can be divided into broad functional groups with different influences on marine geochemistry. Except for the most extreme high temperatures tolerated exclusively by photosynthetic bacteria, there does not appear to be a difference in the range of temperature optima available to different functional groups (Thomas *et al.*, 2012). In contrast, a meta-analysis of published studies examining marine phytoplankton growth rates at modern and future CO_2_ levels revealed a persistent difference between functional groups in terms of the growth rate response (GRR) to elevated CO_2_ (Dutkiewicz *et al.*, 2015). While substantial variability in GRR existed in all functional groups, on average larger organisms (including most eukaryotes) where unaffected by CO_2_ at year 2100 levels, whereas most cyanobacteria (including all tested nitrogen-fixers) grew faster under future conditions. A global ocean mixing model suggested that this discrepancy between functional group CO_2_ GRRs would cause ocean acidification to have a larger impact on the functional composition of future phytoplankton communities than global warming (Dutkiewicz et al., 2015).

Surprisingly, the most far-reaching shift predicted by the model of Dutkiewicz et al. (2015) involved competition between the two most abundant genera of marine cyanobacteria, *Prochlorococcus* and *Synechococcus*, which together contribute more than 50% of fixed carbon in some marine systems (Liu *et al.*, 1997, Campbell *et al.*, 1994). In the modern ocean, *Prochlorococcus* and *Synechococcus* coexist globally but dominate different regions, with *Prochlorococcus* more abundant in permanently stratified oligotrophic waters between 40° N and 40° S latitude, and *Synechococcus* dominating in seasonally mixed and coastal waters (Partensky *et al.*, 1999, Ahlgren & Rocap, 2012). A model of the future ocean considering only temperature increase predicted that both *Prochlorococcus* and *Synechococcus* would increase in abundance, but their ratios would remain roughly similar throughout their range (Flombaum *et al.*, 2013). However, *Prochlorococcus* was the one exception to the rule of increased growth rate at high CO_2_ amongst cyanobacteria (Fu *et al.*, 2007a), and as a consequence the models of Dutkiewicz et al. that considered both temperature increase and ocean acidification predicted that the competitive balance between these two groups would shift sufficiently over the coming century that *Synechococcus* would replace *Prochlorococcus* throughout its range, essentially leading to the worldwide elimination of *Prochlorococcus*’ niche (Dutkiewicz et al., 2015). While this remarkable conclusion was based on a single published experiment with a single cultured strain of each genus (Fu et al., 2007a), subsequent experiments have confirmed that gene expression of the “helper” heterotrophic bacteria that it depends on to tolerate ubiquitous oxidative stress exposure (Morris *et al.*, 2011, Hennon et al., 2018).

Because understanding how critical taxa like *Prochlorococcus* will fare under future CO_2_ conditions is key to our ability to predict how anthropogenic changes will impact the biogeochemistry and biodiversity of future oceans, we sought to increase our understanding of how ocean acidification might influence the interaction between *Prochlorococcus* and *Synechococcus*. First, we measured the GRR of several different isolates from each genus, including several well-characterized ecotypes of each, to determine if the differential responses previously reported (Fu *et al.*, 2007b, Hennon et al., 2018) are representative. Second, we conducted direct, head-to-head competition experiments with *Prochlorococcus* and *Synechococcus* in co-culture to determine if predictions made from unialgal cultures reflected behavior in mixed communities.

## RESULTS

### Growth rate responses to elevated CO_2_ of the two genera

We measured growth rates at modern (∼400 ppm) and year 2100 (∼800 ppm) CO_2_ concentrations for 8 strains of *Synechococcus* and 5 strains of *Prochlorococcus* representing a broad range of the ecotypic diversity in each genus (Table S1). While there was substantial and significant variation at the level of individual strains (linear mixed effects model (LME), p < 0.001), there was not a significant overall difference in exponential growth rates between the genera (LME, p = 0.08), nor was there an effect of CO_2_ treatment on exponential growth rates of either genus (LME, p = 0.42) (Table 1). In contrast, with a single exception all *Prochlorococcus* strains tested exhibited lower realized growth rates (e.g., batch culture growth rates inclusive of lag times) at high CO_2_, whereas *Synechococcus* strains exhibited a broad variety of responses, with a general trend toward enhancement at high CO_2_ (Table 1). As a consequence, *Synechococcus* strains overall had higher realized growth rates than *Prochlorococcus* by 0.14 +/− 0.06 d^−1^ (LME, p = 0.002), and the effect of CO_2_ was significantly negative for *Prochlorococcus* (LME,−0.07 +/− 0.03 d^−1^, p = 0.02) but not for *Synechococcus*.

**Table 1.**
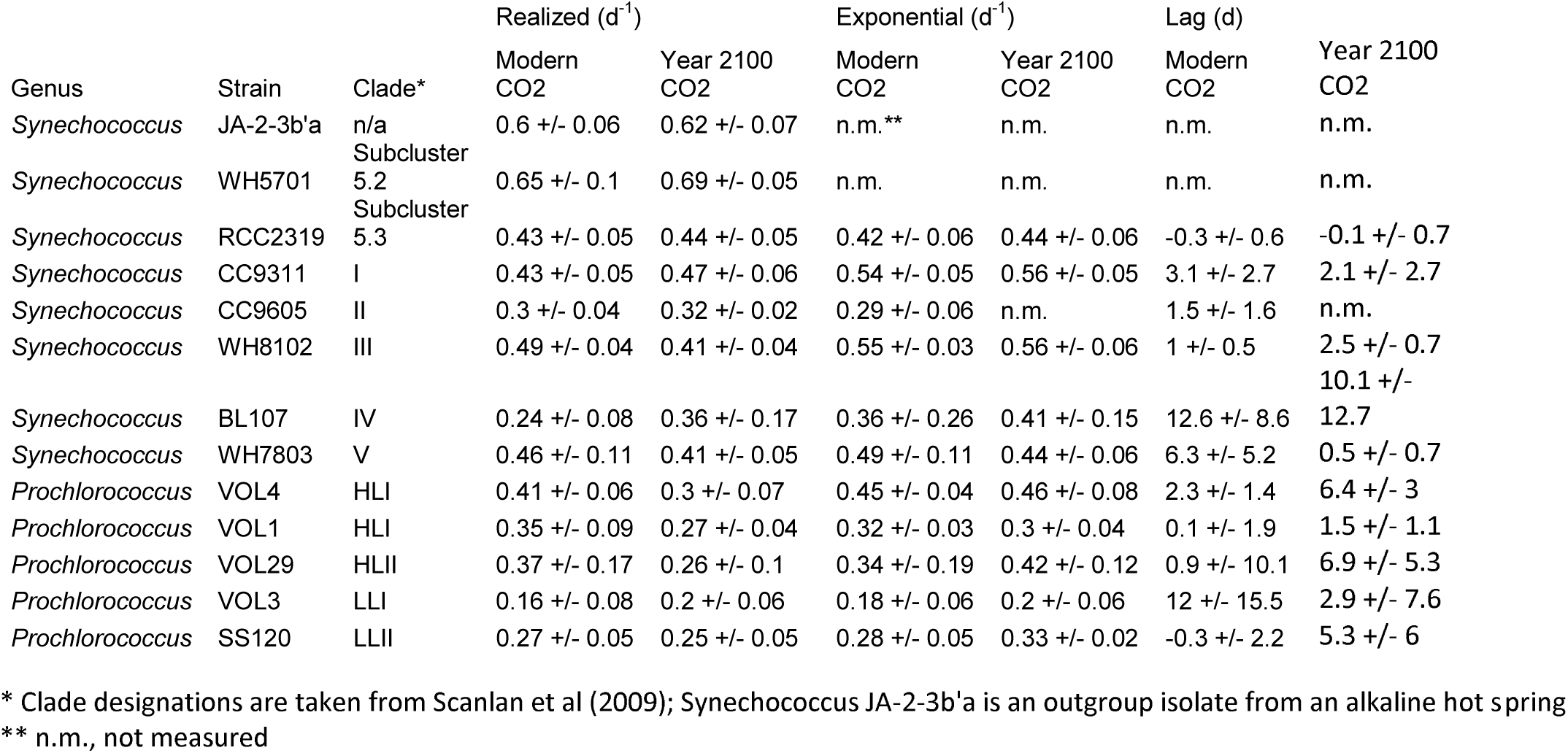
Growth rate parameters for Synechococcus and Prochlorococcus strains. Values are +/− 95% confidence intervals.

We next investigated the GRR to elevated CO_2_ for the genera (Fig. 1). For each strain, we expressed GRR as the ratio of average high CO_2_ growth rate to ambient CO_2_ growth rate; GRRs were calculated separately for exponential and realized growth rate measurements. Consistent with other marine cyanobacteria as revealed by the meta-analysis of Dutkiewicz et al. (2015), we found that the exponential GRR for both genera were similar and significantly greater than 1 (Wilcoxon signed rank test, p = 0.04). The realized GRR, however, showed a starkly opposite trend. Where *Synechococcus*’ realized GRR was nearly identical to its exponential GRR, *Prochlorococcus’* realized GRR was significantly lower than its exponential GRR (LME, p = 0.03). A previous report (Hennon et al., 2018) showed that the discrepancy between *Prochlorococcus*’ exponential and realized growth rates was caused by a longer “lag” after transfer into fresh media, apparently caused by a die-off of cells sometimes exceeding 99% of the total transferred population. Lags were also observed in this experiment (Table 1), and while they were in general longer at high CO_2_ and especially for *Prochlorococcus* at high CO_2_ (by over 5 days) there was sufficient variability between strains in both genera to prevent these effects from being statistically significant.

**Figure 1.**
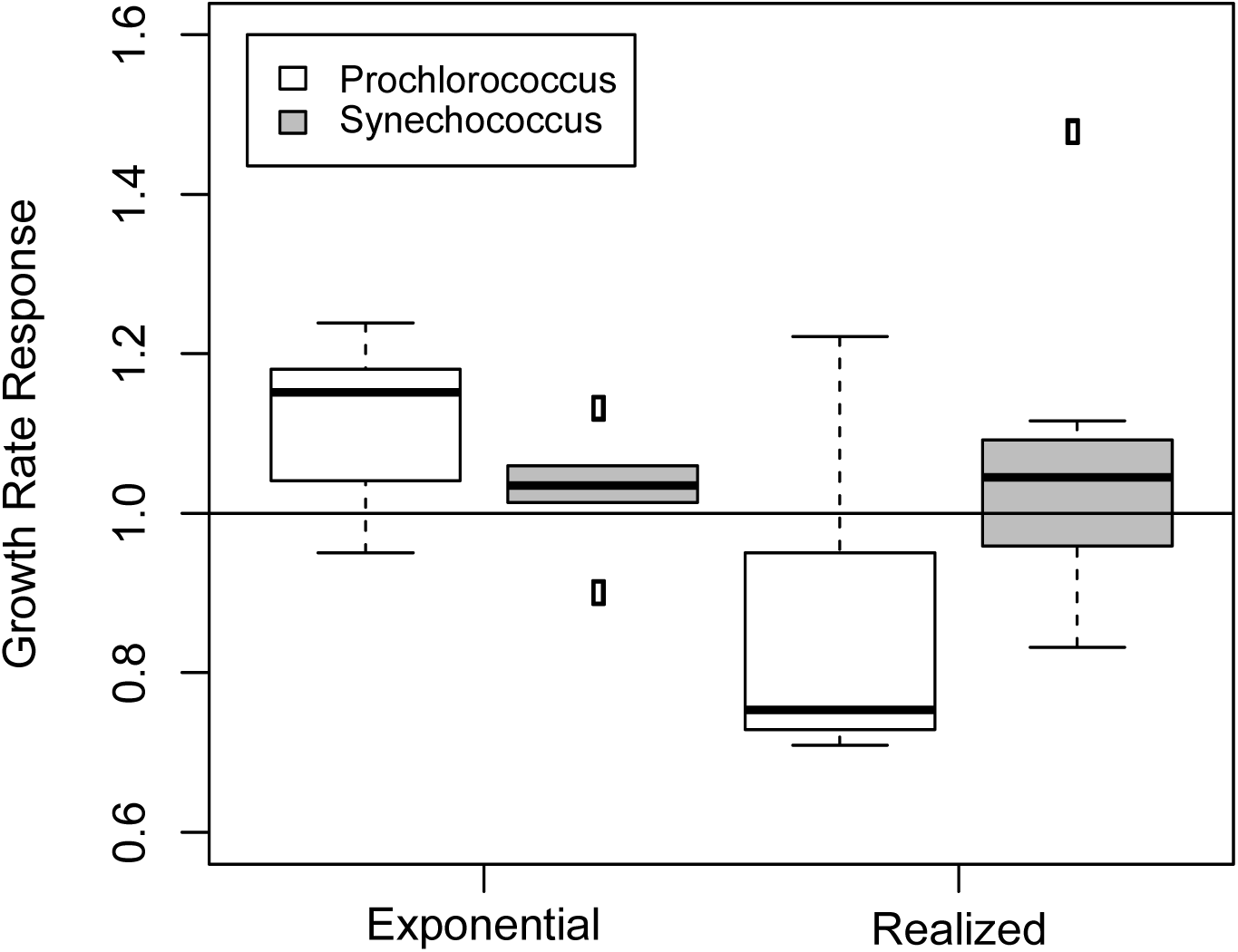
Growth rate responses (GRRs) to CO_2_ for *Prochlorococcus* and *Synechococcus* strains. The ratio of growth rate at year 2100 CO_2_ versus modern CO_2_ was determined for diverse strains of both picocyanobacterial genera. Exponential GRR was calculated using at least 3 data points where cell density was linear with time after log-transformation. Realized (i.e. Malthusian) growth rates were determined based on the time required for a culture to reach its transfer cutoff density, given its initial cell density. Boxes extend from the 25th to 75th percentiles, with the central line representing the median value; whiskers extend to the 5th and 95th percentiles, with values beyond this range shown individually as circles.

### Influence of CO_2_ on the competitive fitness of *Prochlorococcus* vs. *Synechococcus*

Using exclusively GRR estimates from one strain each of *Prochlorococcus* and *Synechococcus* derived from Fu et al. (2007b), Dutkiewicz et al. (2015) predicted that the former would be at a worldwide competitive disadvantage under the predicted year 2100 CO_2_ regime. Our own experiments with unialgal cultures provide conflicting evidence on this question (Fig. 1), with *Prochlorococcus* and *Synechococcus* having similar exponential GRRs, but *Synechococcus* having a sizable advantage in realized GRR. Because microbial fitness can involve a number of components other than maximum exponential growth rate (Vasi *et al.*, 1994), the ideal method to measure relative fitness is through direct competitions in co-culture. Therefore we competed clonal populations of *Prochlorococcus* MIT9312 with *Synechococcus* CC9311 in co-culture under ambient and future CO_2_ conditions. In parallel, we measured the single-culture growth rates of each in the same medium. While none of these organisms were in axenic culture, each was paired with a pure, clonal culture of the same heterotophic bacterium, *Alteromonas* sp. EZ55. Thus, the only variables in these experiments are i) CO_2_ concentration and ii) presence or absence of the competing organism. These strains were chosen because they exhibited opposite GRRs in single culture, and therefore we expected they would provide the clearest test of the efficacy of unialgal culture experiments for predicting competition in mixed communities.

When relative fitness in our first experimental run was calculated as simply the difference in realized growth rates of the two competing organisms, there was no effect of CO_2_ on competitive fitness of *Prochlorococcus*, and the fitness of *Prochlorococcus* in the majority of cultures was lower than that of *Synechococcus* (Fig. 2). However, substantial variation existed in fitness measurements between cultures within each CO_2_ treatment, leading us to examine possible sources of variability. Because competition experiments were conducted over several semi-continuous batch culture transfers, natural selection caused the ratio of *Prochlorococcus* to *Synechococcus* to change over time, leading to the possibility that the relative frequency of the competitors might affect fitness. When we constructed a linear model of *Prochlorococcus* fitness in which the initial frequency of *Prochlorococcus* was a main effect (i.e., a frequency dependence model), the effect of initial *Prochlorococcus* frequency was highly significant but that of the CO_2_ treatment was not (LME. p = 1.29 × 10^−6^ and p = 0.3, respectively, Fig. 3a open circles). Because our initial experiments started at a uniform initial ratio of approximately 10:1 *Prochlorococcus*, we conducted a second series of experiments where we varied the initial ratio from 10:1 to 1:10. These experiments strongly supported the conclusion that *Prochlorococcus’* fitness vs. *Synechococcus* CC9311 exhibits negative frequency dependence with no significant effect of CO_2_ treatment (Fig. 3a, black circles). When both datasets were considered together, *Prochlorococcus* fitness was predicted to decrease by-0.444 +/− 0.092 d^−1^ as its frequency increased from 0 to 100% (LME, p = 2.133 × 10^−8^), but no effect of CO_2_ was detectable (LME, p = 0.13).

**Figure 2.**
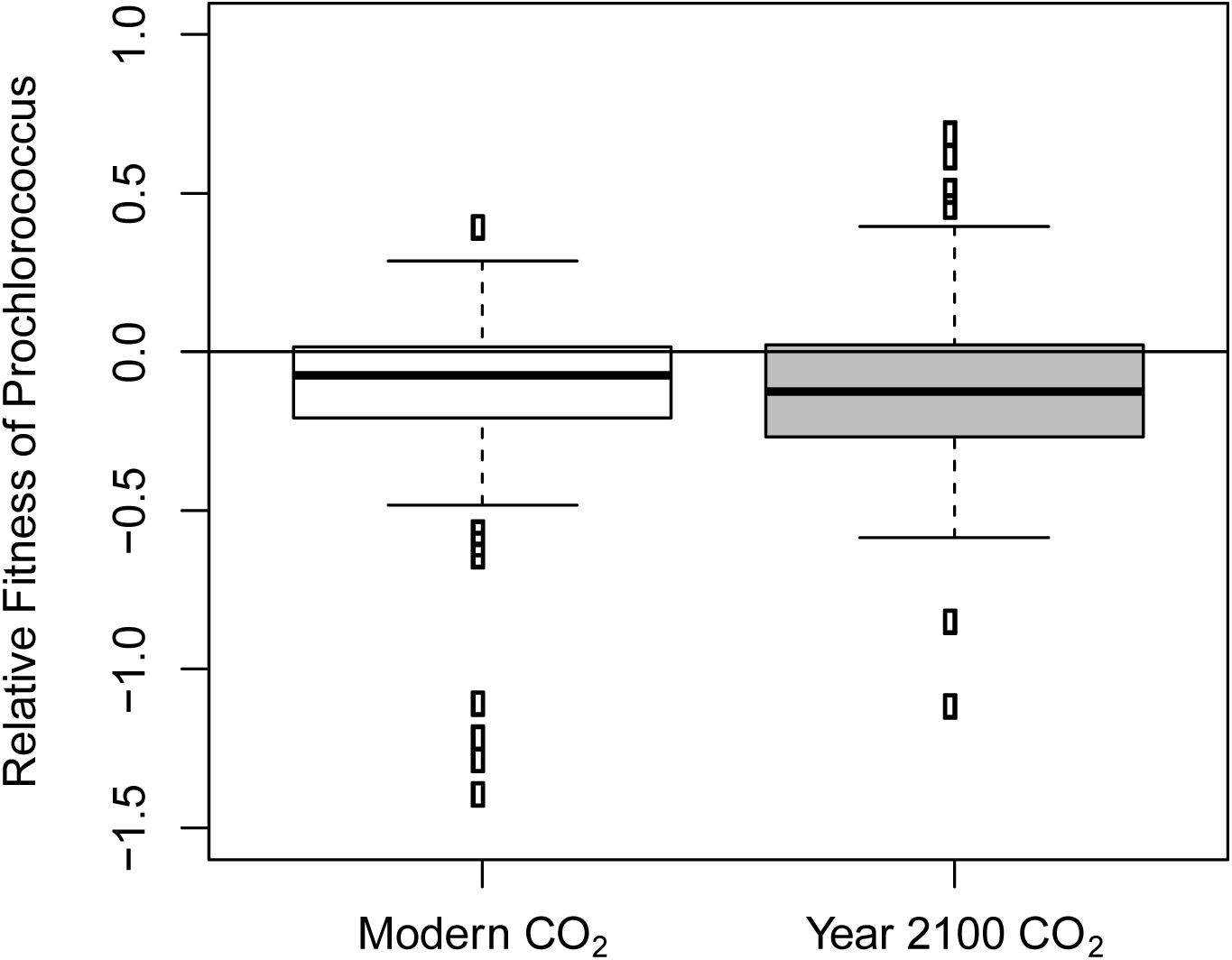
Relative fitness of *Prochlorococcus* versus *Synechococcus* in direct competitions. Fitness was calculated as the *Prochlorococcus* growth rate minus the *Synechococcus* growth rate and is given in units of d^−1^. Boxes extend from the 25th to 75th percentiles, with the central line representing the median value; whiskers extend to the 5th and 95th percentiles, with values beyond this range shown individually as circles.

**Figure 3.**
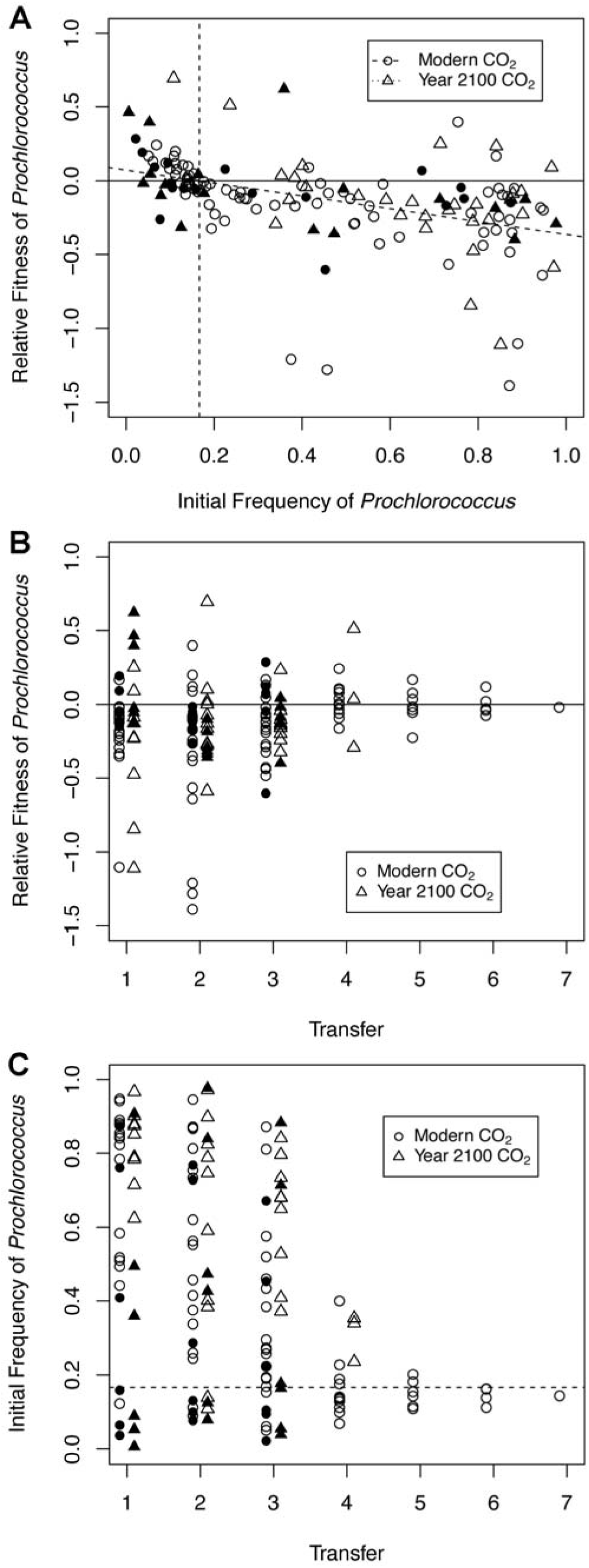
Negative frequency dependence of *Prochlorococcus* fitness. A) The relative fitness of *Prochlorococcus* was strongly frequency dependent, with *Prochlorococcus* tending to have a disadvantage when abundant and an advantage when rare. The broken diagonal line represents the regression of fitness on initial frequency, and the vertical line shows the predicted equilibrium frequency. B) Competition experiments began with *Prochlorococcus* at various ratios with *Synechococcus,* and were continued across at least 3 successive 26-fold transfers into fresh medium. Over time, the relative fitness of *Prochlorococcus* converged on 0, or equal fitness with *Synechococcus*. C) As time progressed in any given experiment, the frequency of *Prochlorococcus* converged on the equilibrium frequency predicted by the regression analysis, shown here with a dotted line. Data from two sets of experiments using similar methods are distinguished here by white and black symbols.

Negative frequency dependence of fitness is one way that very similar organisms can persist in sympatry despite being in competition for a single limiting nutrient. Indeed, in our experiments *Prochlorococcus* generally had greater fitness than *Synechococcus* when initially rare, and lower fitness when initially common (Fig. 3a). The regression of relative fitness on initial frequency (dashed diagonal line, Fig. 3a) suggests that the two organisms should have equal fitness and should be able to coexist when *Prochlorococcus* represents ∼17% of the population (dashed vertical line, Fig. 3a). Supporting this prediction, as competition experiments continued over multiple transfer cycles, the relative fitness of *Prochlorococcus* appeared to converge on neutrality (i.e., equal fitness with *Synechococcus*, Fig. 3b) and the frequency of *Prochlorococcus* approached the prediction of the regression model (Fig. 3c).

### Synergy between *Prochlorococcus* and *Synechococcus*

In the process of conducting the competition experiments, we observed that both MIT9312 and CC9311 appeared to grow faster in co-culture than most strains of either genus grew in unialgal culture. To confirm this, we grew the same MIT9312 and CC9311 clones as unialgal cultures in the same medium and under the same conditions used for the competition experiments. Indeed, we found that the median growth rate of *Prochlorococcus* in the presence of *Synechococcus* was almost twice as fast as in solo culture (LME, p = 1.55 × 10^−5^ for the main effect of strain and 0.0014 for the interaction of strain and co-culture status, Fig. 4). The variability introduced by frequency dependence obscured the effect of co-culture on *Synechococcus* growth rate, but maximum growth rates observed for this strain were also much higher in co-culture. Both organisms approached or exceeded doubling times of 1 d^−1^ in co-culture, consistent with maximum growth rates observed for these organisms in nature (Liu et al., 1997). In this analysis, the effect of CO_2_ on growth rates was only significant as a three-way interaction term with strain and co-culture condition (LME, p = 0.004), indicating that one way that co-culture with *Synechocococcus* CC9311 affects *Prochloroccus* MIT9312 growth is by essentially eliminating the growth defect observed in *Prochlorococcus* unialgal cultures grown at high CO_2_.

**Figure 4.**
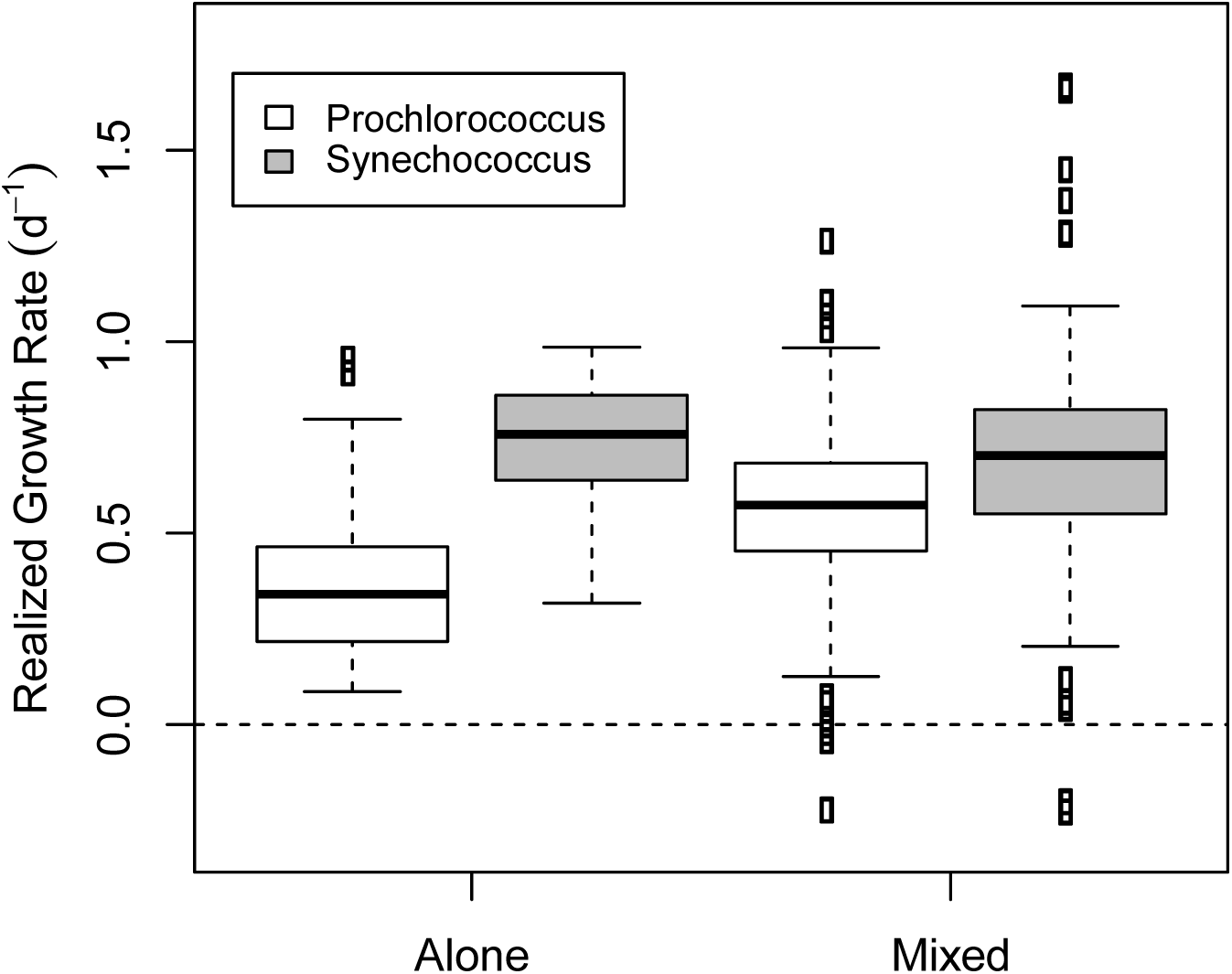
Synergistic interaction between *Prochlorococcus* and *Synechococcus*. The realized (Malthusian) growth rate of *Prochlorococcus* increased significantly in competition co-cultures relative to unialgal culture. While *Synechococcus* growth rates did not increase significantly due to high variability caused by negative frequency dependence, growth rates were observed that were much higher than any measured in unialgal cultures. Boxes extend from the 25th to 75th percentiles, with the central line representing the median value; whiskers extend to the 5th and 95th percentiles, with values beyond this range shown individually as circles.

## DISCUSSION

The results presented here provide useful context for the models of Dutkiewicz et al. (2015) and expand our knowledge about the responses of these key picophytoplankton taxa to changing atmospheric CO_2_. While there was substantial culture-to-culture variability (Table 1), the median exponential GRR of both *Synechococcus* and *Prochlorococcus* was greater than 1 (Fig. 1), consistent with cyanobacterial GRRs previously reported (Endres *et al.*, 2013, Fu et al., 2008, Garcia *et al.*, 2011, Hutchins et al., 2007, Kranz *et al.*, 2010, Fu et al., 2007b). Along with similar elevated GRRs reported for the eukaryotic picophytoplankter *Ostreococcus taurii* (Schaum & Collins, 2014), these observations suggest that very small photoautotrophs may generally benefit from a greater supply of CO_2_ despite not being Liebig limited by access to carbon. A common explanation for the growth enhancing effects of CO_2_ is that cells can reduce their resource investment in carbon concentrating mechanisms (e.g. carboxysomes, bicarbonate transporters) when the CO_2_:HCO_3_^−^ ratio increases at low pH. However, empirical evidence for this speculation remains sparse, and contrary examples exist: larger diatoms, for instance, appear to be more enhanced by CO_2_ than smaller ones (Wu *et al.*, 2014).

*Prochlorococcus* is dependent on the presence of “helper” organisms for survival in stressful conditions (Morris *et al.*, 2008, Morris et al., 2011). We routinely co-culture *Prochlorococcus* with an isolate from the cosmopolitan marine bacterial genus *Alteromonas* to mitigate oxidative stresses *Prochlorococcus* experiences in culture media, but our previous work (Hennon et al., 2018) suggested that elevated CO_2_ altered the interaction between these two organisms in a manner that was ultimately damaging to *Prochlorococcus* resulting in long lag phases and cell death upon culture dilution. Because of this, we predicted that *Prochlorococcus* would be rapidly outcompeted by *Synechococcus* under high CO_2_ conditions, since the latter exhibits no such growth defects. On the contrary, it appears that the presence of *Synechococcus* CC9311 is able to compensate for deficiencies in the helping ability of *Alteromonas* under future CO_2_ concentrations. How it accomplishes this is unclear. The central helping mechanism of *Alteromonas* is the removal of hydrogen peroxide from culture media (Morris et al., 2011), but the *Synechococcus* CC9311 genome does not encode catalase or any other strong hydrogen peroxide scavenging enzymes (Scanlan *et al.*, 2009). Alternatively, it is possible that *Synechococcus* indirectly helps by restoring gene expression in *Alteromonas* to the modern CO_2_ state, or that *Synechococcus* provides help to *Prochlorococcus* through an entirely different mechanism. The fact that *Synechococcus* was able to increase the growth rate of *Prochlorococcus* under both CO_2_ conditions (Fig. 4) strongly suggests that alternative, currently unknown mechanisms of helping exist in these cultures.

While the requirement for help by *Prochlorococcus* is well-established, it was surprising to us that *Synechococcus* CC9311 also appeared to respond positively to co-culture (Fig. 4). Whereas *Prochlorococcus* and *Alteromonas* occupy distinctly separate trophic niches, under our experimental conditions *Prochlorococcus* and *Synechococcus* were in direct scramble competition for all of their nutritional requirements. Assuming that both *Synechococcus* and *Prochlorococcus* were limited by the same nutrient, the competitive exclusion principle (Hardin, 1960) predicts that one or the other should ultimately be driven to extinction. Instead, strong frequency dependence of fitness facilitated their co-existence (Fig. 3). While the current experiments do not reveal what mechanism facilitates this mutual helping interaction, this pattern of negative frequency dependence is consistent with the predictions of the Black Queen Hypothesis (Morris *et al.*, 2012, Morris, 2015), which describes an evolutionary path where selection for loss of genes that produce publically-available “leaky” products can favor interdependency in microbial communities. It is thought that *Prochlorococcus*’ dependence on helpers evolved because other members of its community were unable to keep from removing hydrogen peroxide from the environment at the same time they detoxified their own cytoplasm, providing an opportunity for *Prochlorococcus’* ancestors to lose their catalase gene and gain a growth advantage. This Black Queen process of leaky function loss appears to have broadly affected marine bacteria and has led to extensive networks of interdependence in oceanic microbial communities. It is possible that CC9311 (and possibly other *Synechococcus* strains as well) have also evolutionarily lost functions that render them dependent on co-occurring microbes, and that *Prochlorococcus* can in some cases act as a helper to these taxa.

Several factors should be considered when relating our experimental results to the ecological interactions between *Prochlorococcus* and *Synechococcus* in the ocean. First, the particular strains used for the competition experiments were chosen because of their relative ease of use in the laboratory and their opposite responses to elevated CO_2_ (Table 1), but they are not commonly found together in nature. *Prochlorococcus* MIT9312 is a tropical, open ocean strain, and *Synechococcus* CC9311 is a temperate coastal strain. Nevertheless, their ranges do overlap (for instance, in the Sargasso Sea (Ahlgren & Rocap, 2012)) and they share many ecological and metabolic characteristics with other members of the same genera that do routinely coexist in the ocean (Scanlan et al., 2009). While it is possible that the negative frequency dependence observed here is a peculiar and fortuitous property of this particular strain pairing, we find that explanation highly unlikely. Second, a substantial fraction of the change in *Prochlorococcus’* growth characteristics between unialgal culture and co-culture with *Synechococcus* can be attributed to a reduction in the lag experienced on transfer to new, elevated CO_2_ media. Lag phase is a property of laboratory batch culture and is arguably not relevant to natural systems, particularly highly stable steady-state systems like the pelagic ocean. However, the factors that contribute to lag phase -- changing environmental conditions, and in particular for *Prochlorococcus* exposure to untreated media containing hydrogen peroxide -- do occur in the ocean, despite its relative environmental stability. For instance, the hydrogen peroxide concentration of the surface ocean can vary by a factor of 4 over a 24 h cycle (Morris *et al.*, 2016), and wind-forced mixing can move cells rapidly from high-intensity light zones near the surface to much dimmer regions at the base of the mixed layer. In addition to its lower contingent of oxidative stress resistance genes, *Prochlorococcus* is also deficient in genes for sensing its environment (Scanlan et al., 2009), and may depend on diverse members of its microbial community to navigate these changes much as it appears to require help to tolerate exposure to new culture media.

Given the above caveats, two important ecological conclusions can be drawn from our experiments. First, attempts to use observations of species in isolation to predict how they will fare in a changing environment should be approached with skepticism. Depending on how the experiments were performed and how growth rate was measured, measurements of CO_2_ GRR in unialgal cultures of *Prochlorococcus* and *Synechococcus* could encourage the mistaken conclusion that *Synechococcus* would have much higher competitive fitness under elevated CO_2_ than *Prochlorococcus.* In the case of the strains studied here, the absolute growth rates of both organisms would also be significantly underestimated using unialgal cultures. Other recent studies have also demonstrated that phytoplankton responses to CO_2_ can be very different in co-culture, in some cases even reversing the sign of the response completely (Sampaio *et al.*, 2017). To make matters worse, the key finding of this experiment -- negative frequency dependence between the two strains -- would have escaped our notice if we had not followed the fates of these cultures through multiple transfers with fluctuating starting densities. Negative frequency dependence-stabilized polymorphisms arise readily during microbial evolution and can persist for at least tens of thousands of generations (Rozen & Lenski, 2000, Elena & Lenski, 1997, Good *et al.*, 2017) but are easily missed using common methods for measuring fitness {Ribeck, 2015 #4523}. Moreover, Black Queen and other frequency dependent processes are likely to be very influential in the evolutionary fates of microbes in the ocean and other natural environments (Cordero & Polz, 2014, Braakman *et al.*, 2017). Thus, the importance of both direct competition experiments and frequency-dependent experiments for predicting the fates of changing microbial ecosystems cannot be overstated.

Second, our results provide an alternative hypothesis for the widespread coexistence of *Prochlorococcus* and *Synechococcus* in the world’s oceans. Both strains coexist across the entire temperate and tropical open ocean; for example, across an Atlantic Meridional Transect sampling cruise from 50° N to 50° S, *Prochlorococcus*:*Synechococcus* ratios ranged from 1:10 to 100:1, but both types were constantly present (Johnson *et al.*, 2006). Coexistence of these strains is often explained by their divergent nitrogen requirements: *Synechococcus* can utilize nitrate-N, whereas *Prochlorococcus* generally cannot (Bragg *et al.*, 2010). However, nitrate availability was not a strong predictor of *Synechococcus* abundance globally (Flombaum et al., 2013), and nitrate-utilizing strains of *Prochlorococcus* are known {Berube, 2015 #4081;Martiny, 2009 #5185}. Another possibility is that the coexistence of these two taxa is an example of the classic “Paradox of the Plankton” (Hutchinson, 1961), where many species with nearly identical niches are thought to be able to coexist because of the relatively more rapid rate of environmental change compared to competitive exclusion (e.g. Chandler *et al.*, 2016). Our experiments provide a third possibility: that these similar taxa, like many other microorganisms in their habitat, have evolved dependencies on one another that prevent either from permanently dominating their shared nutritional/environmental niche. While the coexistence frequency predicted from our competition experiments (X-intercept of the regression line in Fig. 3a) are products of a particular culture environment and are not directly relatable to any natural ecosystem, we can nevertheless conclude that the same dynamics that allow coexistence in our simplified cultures could also contribute to these and related strains coexisting within the ratios observed in the actual ocean.

In conclusion, we have shown that the exponential growth rates of diverse strains of *Prochlorococcus* and *Synechococcus* respond to high CO_2_ in a manner consistent with other cyanobacteria and small phytoplankton, although *Prochlorococcus* manifests growth defects at high CO_2_ apparently related to its requirements for “help” from other members of the community. We have also shown that co-cultures of *Prochlorococcus* and *Synechococcus* behave much differently than we would expect from cultures with only one or the other, suggesting both that predictions from single cultures should be met with skepticism, and also that two-way, mutualistic interactions between these taxa may contribute to their coexistence in the ocean. Finally, our results highlight the need for a better understanding of how interactions within the whole community control the relative fitness of each component species, especially for systems such as the ocean where accurate predictions of future states are so critical.

## METHODS

### Strains and culture conditions

The strains used in this experiment are listed in Supplemental Table 1. All cultures were grown in a Percival growth chamber under cool fluorescent lights at 21° C under 150 μmol photons m^−2^ s^−1^ on a 14:10 light:dark cycle. Cultures were grown in 13 mL of medium in sterilized, acid-washed, conical-bottom, screw-cap glass centrifuge tubes. All experiments with *Synechococcus* strains, including co-culture experiments, were incubated on a rotating tissue culture wheel (Thermo) at approximately 60 rpm. Because its small size prevents *Prochlorococcus* from settling over relatively short time scales, experiments containing only *Prochlorococcus* were conducted in static test tube racks under identical environmental and light conditions. Culture media were prepared in an artificial seawater (ASW) base (per 20 L: 568.22 g NaCl, 15.78 g KCl, 31.58g CaCl_2_ * 2H_2_O, 144.28 g MgSO_4_ * 7H_2_O, 103.6 g MgCl2 * 6H2O) that was autoclaved in 2 L batches in acid-washed glass bottles. After cooling, filter-sterilized NaHCO_3_ was added to a final concentration of approximately 2.3 mM. The resulting mixture was bubbled with sterile filtered air from an aquarium pump overnight to equilibrate with the ambient atmosphere. ASW was amended with nutrient stocks to produce finished media (see Andersen (2005) for trace metal and vitamin stock recipes). *Synechococcus-*only cultures were grown in SEv medium (32 μM NaNO_3_, 2 μM NaH_2_PO_4_, 20 μL L^−1^ SN trace metal stock, and 20 μL L^−1^ F/2 vitamin stock). *Prochlorococcus*-only cultures were grown in PEv medium (32 μM NH_4_Cl, 2 μM NaH_2_PO_4_, and 40 μL L^−1^ Pro99 trace metal stock). Mixed competition cultures were grown in PEv supplemented with 20 μL L^−1^ F/2 vitamin stock. Base carbonate parameters of the finished medium were assayed once prior to use (see below). Prior to inoculation, CO_2_ concentrations were manipulated by addition of standardized stocks of either hydrochloric acid and bicarbonate (to achieve elevated CO_2_ conditions) or, if necessary, sodium hydroxide (to compensate for high ambient indoor CO_2_ levels in the laboratory). To minimize pH shift during a growth experiment, cultures were grown with minimal headspace, caps were sealed airtight, and daily sampling was performed quickly. Additionally, cultures were diluted into fresh media before growing to sufficient density for their carbon concentrating mechanisms to affect pH.

### Growth rate and competition experiments

Prior to experiments, *Prochlorococcus* and *Synechococcus* cultures were acclimated to the experimental culture medium under either ambient (∼400 ppm) or year 2100 CO_2_ (∼800 ppm) conditions for approximately 5 generations (i.e., one transfer cycle). For competition cultures, *Prochlorococcus* and *Synechococcus* were acclimated to competition conditions as unialgal cultures before mixing into competition cultures. For unialgal growth rate experiments and the initial competition experiments, initial cell densities were set to approximately 10^4^ cells mL^−1^ *Synechococcus* or 10^5^ cells mL^−1^ *Prochlorococcus*. This 10:1 ratio was chosen because *Prochlorococcus* cells are substantially smaller than *Synechococcus* cells, so the divergent ratio led to a more equitable starting concentration in terms of total carbon/chlorophyll. Subsequent competition experiments varied this initial ratio by leaving the initial *Synechococcus* concentration at a constant 10^4^ cells mL^−1^ and titrating *Prochlorococcus* appropriately. Culture tubes were assayed for cell concentrations approximately every two days. 10 μL aliquots were diluted tenfold into sterile ASW in a microtiter plate and then examined using a Guava HT flow cytometer (Millipore). Cell types were differentiated based on forward scattering of blue laser light and intensity of red chlorophyll autofluorescence. Using these parameters, *Prochlorococcus* and *Synechococcus* populations were readily distinguished from each other. Whenever cell density crossed a cutoff (2.6 × 10^6^ cells mL^−1^ for *Prochlorococcus*, 2.6 × 10^5^ cells mL^−1^ for *Synechococcus*), the culture was diluted 26-fold into fresh media set to the same CO_2_ condition. For competition cultures, transfers were performed whenever either competitor crossed its cutoff cell density. All experiments were run for at least 3 transfer cycles, or approximately 14 generations. The realized growth rates, exponential growth rates, and lag times of the phytoplankton cultures were calculated as described in Hennon et al. (2018).

### Chemical Analysis of Media

The carbonate parameters of our media were calculated using pH and alkalinity measurements. A Mettler-Toledo automatic titrator was used for all alkalinity assays. ASW pH was determined by comparison with standardized Tris and 2-aminopyridine buffers using a Mettler-Toledo pH probe or by using *m*-cresol purple dye (Dickson et al., 2007). Titrations were performed automatically using a 0.1N HCl standard solution purchased from Fisher and not standardized further. Additionally, salinity was determined using a salinity pen (Sper Scientific), and density was determined by weighing a 35 mL aliquot using a Mettler-Toledo scale. Carbonate parameters of the media used for the experiments reported here are given in Supplemental Table 2.

To determine the volumes of HCl / HCO_3_^−^ or NaOH additions needed to mimic the oceanic carbonate system for the present day or projected year 2100, we used the *seacarb* package in R (Gattuso & Lavigne, 2009). After determination of the baseline carbonate chemistry (i.e., pH and alkalinity) of a batch of medium, the *oa* function in *seacarb* was used to determine additions of calibrated sodium bicarbonate and hydrochloric acid stocks necessary to adjust pCO_2_ to either 400 ppm or 800 ppm. Stocks used for CO_2_ manipulation were carefully calibrated by measuring how aliquots altered the alkalinity of an ASW sample; triplicate titrations were performed to calibrate each standard solution.

### Statistical analyses

All statistics were calculated in R v 3.3.1. Unless otherwise indicated, all statistics mentioned in the text are derived from linear mixed effects models computed using the *lme4* package. Fixed effects were one or more of the following: phytoplankton genus, CO_2_ treatment, and whether a culture was unialgal or a mixed competition co-culture. Random effects were the individual strains tested, either specific clone pairings (for the competition experiments) or different species / strains (for the growth rate / GRR experiments). Significance levels for the fixed effects were determined by running the full model, then running the model with each fixed effect eliminated, then comparing the two model fits with an *F* test. The complete R code and raw data are available with the supplemental information and via download from BCO-DMO.

In the competition experiments, a single outlier data point (showing a fitness value approximately twice the next highest value) was removed prior to analysis. This outlier was likely the result of a mistranscribed exponent in a cell density measurement; since its veracity could not be confirmed, we eliminated the point. None of the conclusions of this work were changed by the exclusion of this outlier.

## ACKNOWLEDGEMENTS

Strains used in this study were generously provided by Steve Wilhelm, Erik Zinser, and Jeff Krause. We acknowledge the assistance of Alex Durrant, Elizabeth Entwistle, and Christina Cooley for laboratory assistance and Gwenn Hennon and Erik Zinser for valuable comments on an early draft of this manuscript. We thank the faculty and staff of the Alabama School of Fine Arts for supporting Margaret Knight’s research on this project, and Charles Amsler for suggesting her collaboration with the Morris research groups. This work was supported by NSF grant OCE-1540158 to JJM.

---

**Supplementary Table S1.** Strains used in this study.

| **Strain** | **Alias** | **Genus** | **Clade\*\*** | **Origin** |
| --- | --- | --- | --- | --- |
| JA-2-3b'a | n/a | *Synechococcus* | n/a | Steven Wilhelm |
| WH5701 | CCMP1333 | *Synechococcus* | Subcluster 5.2 | Jeff Krause |
| RCC2319 | n/a | *Synechococcus* | Subcluster 5.3 | RCC*** |
| CC9311 | CCMP2515 | *Synechococcus* | I | Jeff Krause |
| CC9605 | RCC753 | *Synechococcus* | II | RCC |
| WH8102 | CCMP2370 | *Synechococcus* | III | NCMA**** |
| BL107 | RCC515 | *Synechococcus* | IV | RCC |
| WH7803 | CCMP1334 | *Synechococcus* | V | Jeff Krause |
| MIT9312* | VOL4 | *Prochlorococcus* | HLI | Erik Zinser |
| MIT9215* | VOL1 | *Prochlorococcus* | HLI | Erik Zinser |
| VOL29* | n/a | *Prochlorococcus* | HLII | Erik Zinser |
| NATL2A* | VOL3 | *Prochlorococcus* | LLI | Erik Zinser |
| SS120 | n/a | *Prochlorococcus* | LLII | Erik Zinser |

\* These strains were selected for streptomycin resistance as described in Morris et al. (2008) and rendered axenic as described in Morris et al (2011) prior to being re-inoculated with pure cultures of the helper *Alteromonas* EZ55
\*\* Clade designations are taken from Scanlan et al (2009); *Synechococcus* JA-2-3b'a is an isolate from an alkaline hot spring
\*\*\* RCC, Roscoff Culture Collection, Station Biologique, France
\*\*\*\* NCMA, National Center for Marine Algae and Microbiota, Bigelow, ME

**Supplementary Table S2.** Carbonate parameters of culture media. Values are means ± 95% confidence intervals for all of the cultures from the *Prochlorococcus* and *Synechococcus* growth curve experiments, as well as the competition experiments.

|  | Present Day |  |  |  | Year 2100 |  |  |  |
| --- | --- | --- | --- | --- | --- | --- | --- | --- |
|  | n | alkalinity<br>(uM CO<sub>3</sub>) | pH | pCO<sub>2</sub><br>(ppm) | n | alkalinity<br>(uM CO<sub>3</sub>) | pH | pCO<sub>2</sub> |
| *Prochlorococcus* | 82 | 2351±5 | 8.049±0.016 | 416±18 | 68 | 2299±4 | 7.730±0.001 | 939±3 |
| *Synechococcus* | 178 | 2325±15 | 8.029±0.008 | 434±10 | 137 | 2294±13 | 7.761±0.005 | 878±11 |
| Competitions | 96 | 2412±5 | 8.085±0.001 | 381±2 | 49 | 2385±15 | 7.781±0.01 | 860±17 |

